# Diversity and composition of methanotroph communities in caves

**DOI:** 10.1101/412213

**Authors:** Kevin D. Webster, Arndt Schimmelmann, Agnieszka Drobniak, Maria Mastalerz, Laura Rosales Lagarde, Penelope J. Boston, Jay T. Lennon

## Abstract

Methane oxidizing microorganisms (methanotrophs) are ubiquitous in the environment and represent a major sink for the greenhouse gas methane (CH_4_). Recent studies have demonstrated that methanotrophs are abundant and contribute to CH_4_ dynamics in caves. However, very little is known about what controls the distribution and abundance of methanotrophs in subterranean ecosystems. Here, we report a survey of soils collected from > 20 caves in North America to elucidate the factors shaping cave methanotroph communities. Using 16S rRNA sequencing, we recovered methanotrophs from nearly all (98 %) of the samples, including cave sites where CH_4_ concentrations were at or below detection limits (≤ 0.3 ppmv). We identified a core methanotroph community among caves that was comprised of high-affinity methanotrophs. Although associated with local-scale mineralogy, methanotroph composition did not systematically vary between the entrances and interior of caves, where CH_4_ concentrations varied. We also observed that methanotrophs are able to disperse readily between cave systems showing these organisms have low barriers to dispersal. Last, the relative abundance of methanotrophs was positively correlated with cave-air CH_4_ concentrations suggesting that these microorganisms contribute to CH_4_ flux in subterranean ecosystems.

**IMPORTANCE:** Recent observations have shown that the atmospheric greenhouse gas methane (CH_4_) is consumed by microorganisms (methanotrophs) in caves at rates comparable to CH_4_ oxidation in surface soils. Caves are abundant in karst landscapes that comprise 14 % of Earth’s land surface area, and therefore may represent a potentially important, but overlooked CH_4_ sink. We sampled cave soils to gain a better understand the community composition and structure of cave methanotrophs. Our results show that the members of the USC-*γ* clade are dominant in cave communities and can easily disperse through the environment, that methanotroph relative abundance was correlated with local scale mineralogy of soils, and that the relative abundance of methanotrophs was positively correlated with CH_4_ concentrations in cave air.

## INTRODUCTION

Methane (CH_4_) is as major greenhouse gas in Earth’s atmosphere, and an energy source for humanity (1, 2). Because of this, it is important to identify the pathways that produce and consume CH_4_. Two major processes are responsible for the removal of CH_4_ from Earth’s atmosphere, oxidation by atmospheric hydroxyl radicals, and by methane-consuming microorganisms (methanotrophs) in surface soils and waters. Top-down and bottom up estimates of yearly atmospheric CH_4_ removal indicate hydroxyl radicals account for 90 % of the sink while methanotrophs consume roughly 5 % (3). Despite the influence that methanotrophs have on regulating Earth’s climate, gaps remain in our understanding of methanotrophs, like how they respond to changing CH_4_ concentrations or how they disperse through the environment (4, 5).

Rocks that host caves are widespread on Earth, and are emerging as an environment that consumes atmospheric CH_4_ (6–11) (Figure 1). Recent studies of cave microbial communities have revealed the presence of atmospheric-CH_4_ consuming methanotrophs in caves (8, 12–15). The methanotrophs that are thought to be responsible for this are termed high-affinity methanotrophs, and have been previously observed from upland soils. These groups are referred to as the Upland Soil Cluster (USC) -*α* and -*γ* clades. Intermittent or unsustained atmospheric CH_4_ oxidation has also been observed in methanotrophs among the families of the Alphaproteobacteria (the *Methylocystaceae* and *Beijernickiaceae*), but it is generally thought that these organisms contribute little to atmospheric CH_4_ consumption (16). Both high-affinity and low-affinity alphaproteobacterial methanotrophs tend to be the most prominent members of the methanotrophic community in environments with low CH_4_ concentrations and high O_2_ concentrations. They are less common in environments with high CH_4_ and low O_2_ concentrations, like lakes or geological seeps, which are dominated by low-affinity methanotrophs found among families belonging to the Gammaproteobacteria (Methylococcaceae and Methylothermaceae) (17, 18).

**Figure 1:**
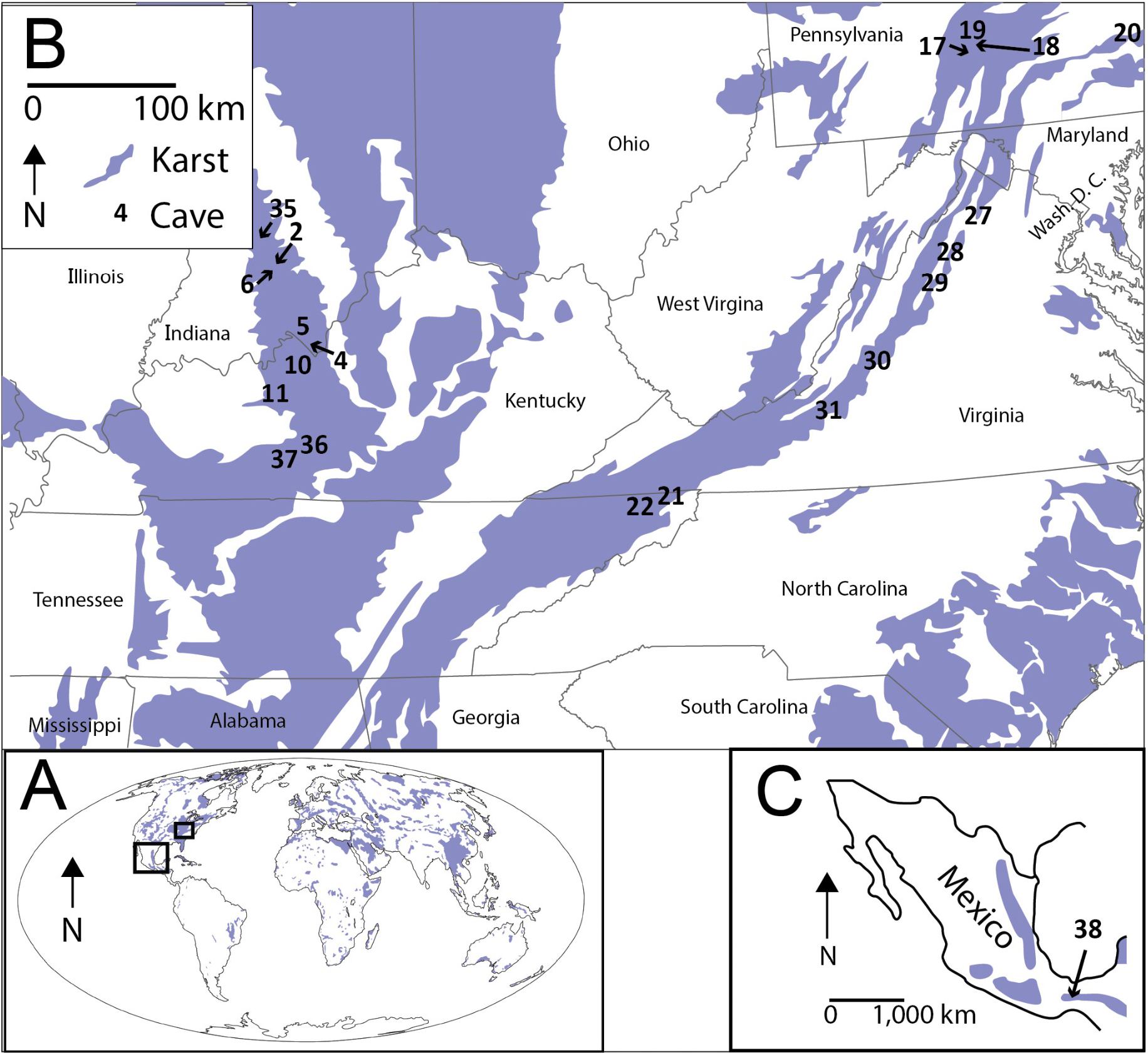
(A) The distributions of karst landscapes at global scale. Inserts show the occurrence of karst (B) in the eastern United States (C) and in Mexico. Numbers represent the locations of sampled caves in this study. Karst land cover data were obtained from (6, 58).

The factors that control the distribution and abundance of methanotrophs in caves are not well understood. Previous work has shown that methanotrophs are frequently related to CH_4_ concentrations, soil texture, and the abundances of other microorganisms (19), but it is unclear how these factors, or the local mineralogy influences methanotroph communities in caves. For example, low-affinity methanotrophs exhibit niche partitioning between environments with high and low CH_4_ concentrations (18, 20). However, it is unknown if methanotrophic communities exhibit similar changes in community composition along atmospheric to subatmospheric gradients of CH_4_.

In addition, very little is known about the spatial distribution of methanotrophs, even though such information could shed light on the biogeography of microbes within and among cave ecosystems. While the community composition of methanotrophs may be influenced by environmental conditions of caves, their abundances may also be influenced by their capacity to disperse. Caves are unique and insular habitats that have a patchy distribution in the terrestrial landscape. However, cells may move through cracks and pores from surface environments or aeolian transport into nearby cave systems. These potentially stochastic modes of dispersal could explain be important in structuring cave methanotroph communities and may help explain the global distribution of methanotrophs. For example, similar methanotrophs have been observed in both Hawai’i and the Arctic (4) suggesting that the movement of these organisms may be relatively unhindered. However, biota in caves are typically unique due to their isolation from other environments (21, 22). Thus, caves provide an ideal environment to test whether methanotrophs are able to disperse readily across the environment or are limited by their ability to disperse.

In this study, we sampled 42 cave soils from 21 limestone caves in North America to examine the factors regulating the community composition of cave methanotrophs (Figure 1). First, we hypothesized that the methanotroph composition should be similar to what has been reported for methanotrophic communities in other low CH_4_ environments, and that the relative abundance of methanotrophs should be correlated with cave-air CH_4_ concentrations. Additionally, we hypothesized that, on the local scale, methanotrophs would be affected by the soil texture and mineral composition of cave soils since these features influence the diffusion of air and micronutrient abundances. Finally, on the regional scale we hypothesized that high-affinity methanotrophs should be able to disperse easily between caves.

## RESULTS

### Methanotroph community structure

Methanotrophs were present in nearly all caves. Based on 16s rRNA sequencing we identified methanotrophs in 98 % of the cave-soil samples, including locations where CH_4_ concentrations were at or below the analytical detection limit (≤ 0.3 ppmv). The maximum relative abundance of methanotrophs in the cave soil samples was 6.25 % and the median relative abundance was 0.88 %. The average site contained 6 OTUs belonging to methanotrophs. High-affinity methanotrophs comprised a median of 99.0 % of the total methanotrophic community at locations where methanotrophs were observed. Members of the USC-*γ* clade represented 47.7 % of the community on average (median), while the members of the USC-*α* clade represented 43.1 % of the community on average (median). Low-affinity methanotrophs represented 0.09 % of the observed methanotroph community on average (median). Members of the USC-*γ* and -*α* were represented by 22 and nine OTUs, repectively, whereas low-affinity methanotrophs belonging to the Alphaproteobacteria and Gammaproteobacteria were represented by 18 and 22 unique OTUs, respectively. Members of the USC-*γ* clade were dominant in the sampled caves (Figure 2). The members of methanotrophic community from Cave 38 (Mexico) were solely high-affinity methanotophs.

**Figure 2:**
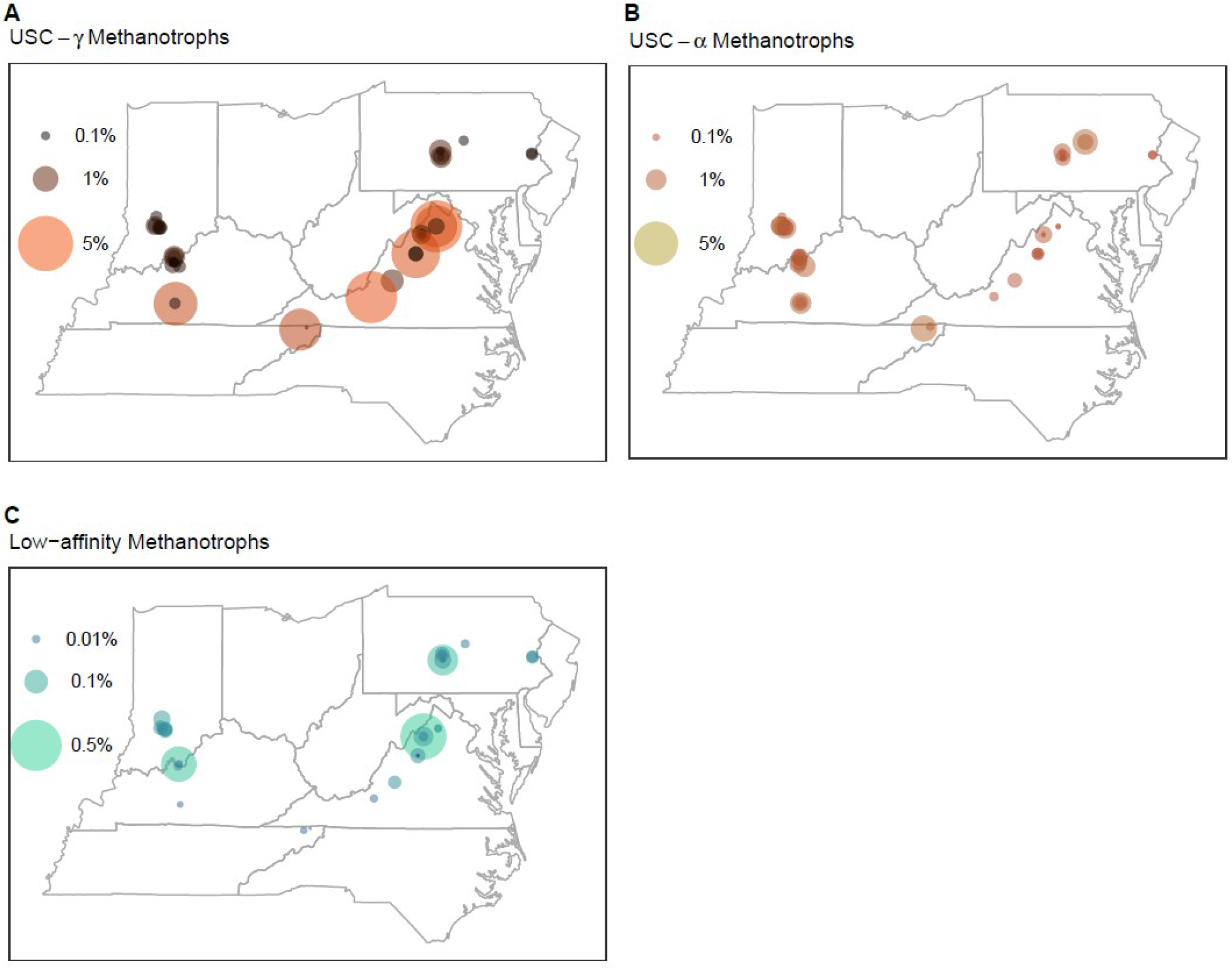
The relative abundance of A) USC-*γ* methanotrophs, B) USC-*α* methanotrophs, and C) Low-affinity methanotrophs in the sampled caves.

### Core Methanotroph Community

We defined taxa that were present in at least 60 % of samples as the core methanotroph community (23, 24). Based on this, the core methanotroph community was made up of three OTUs. Two of these OTUs were from members of the USC-*α* and one was from the USC-*γ*. These taxa were found together in 79 % of samples, and the most widely distributed OTU was observed in 93 % of samples. The core methanotroph community accounted for 97 % of the total methanotroph sequences recovered across all cave samples.

### Environmental influences on community structure

Methanotroph composition was related to the mineralogy of cave soils (Figure 3). The total methanotrophic community was related to the abundance of clinochlorite (multiple regression, *P* = 0.008), muscovite (multiple regression, *P* = 0.03), and microcline (multiple regression, *P* = 0.007). High-affinity methanotrophs showed relationships with the abundance of clinochlorite (multiple regression, *P* = 0.03), and microcline (multiple regression, *P* = 0.02), while the low-affinity methanotrophs were not related to the abundance of any observed mineral (*P* > 0.05). In contrast to mineralogical associations, the composition of the methanotrophic community was not related to the distributions of grain size that comprised a soil (regression, *P >* 0.05).

**Figure 3:**
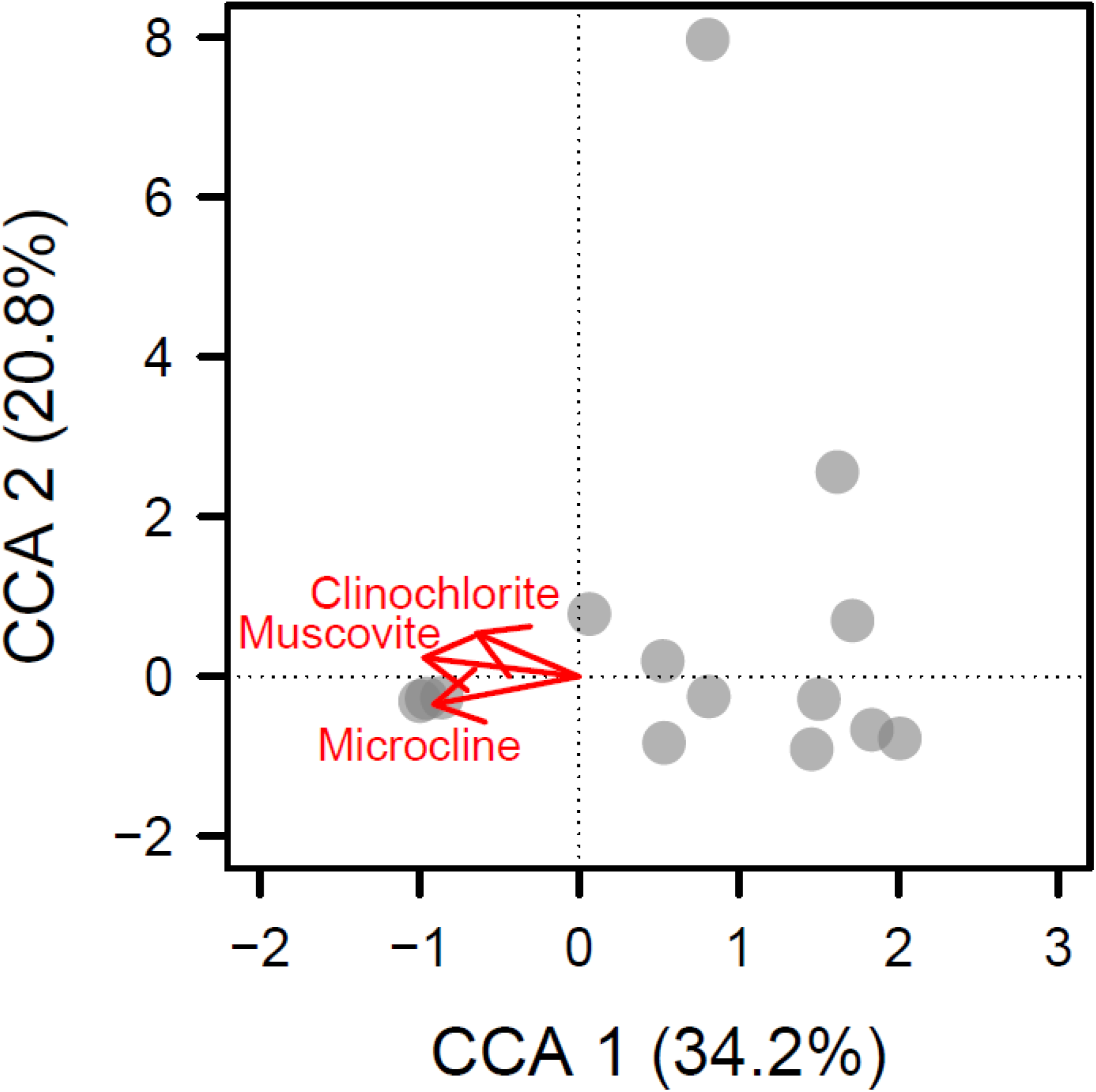
The total methanotrophic community showed relationships with the abundance of muscovite, clinochlorite, and microcline detected in the samples. The vectors show increasing abundances of minerals the samples.

### Biogeography

In ecology, the distance decay relationship (DDR) describes how community similarity changes with increasing geographic distance and environmental similarity (25, 26). Briefly, as the space between two points increases, both biological communities and the environment tend to become less similar. The degree to which species can disperse through the environment can alter how quickly the community assemblage changes across space. Communities that are hindered by dispersal tend to change more quickly than measures of environmental similarity, while communities that have have a high capacity to disperse tend to change as quickly as measures of environmental similarity. Within the methanotroph assemblages, we found that the composition of high-affinity taxa changed at the same rate as geographic distance and measures of environmental similarity (*m*_*high-afinnity*_ = -0.09; *m*_*env*_ = -0.05; Difference of slopes test, *P* = 0.10) (Figure 4). Meanwhile, the composition of low-affinity methanotrophs and the measures of environmental similarity changed little with geographic distance (*m*_*low-affinity*_ = 0.10 ± 0.06; Linear regression, *P* = 0.09; *m*_*env*_ = −0.04 ± 0.05; Linear regression, *P* = 0.43). At smaller spatial scales, the relative abundance of methanotrophs was not related to the distance from a cave entrance (High-affinity: Spearman’s rank correlations, *ρ* = −0.15, S = 4195, *P* = 0.45; Low-affinity: Spearman’s rank correlations, *ρ* = 0.31, S = 2488, *P* = 0.09).

**Figure 4:**
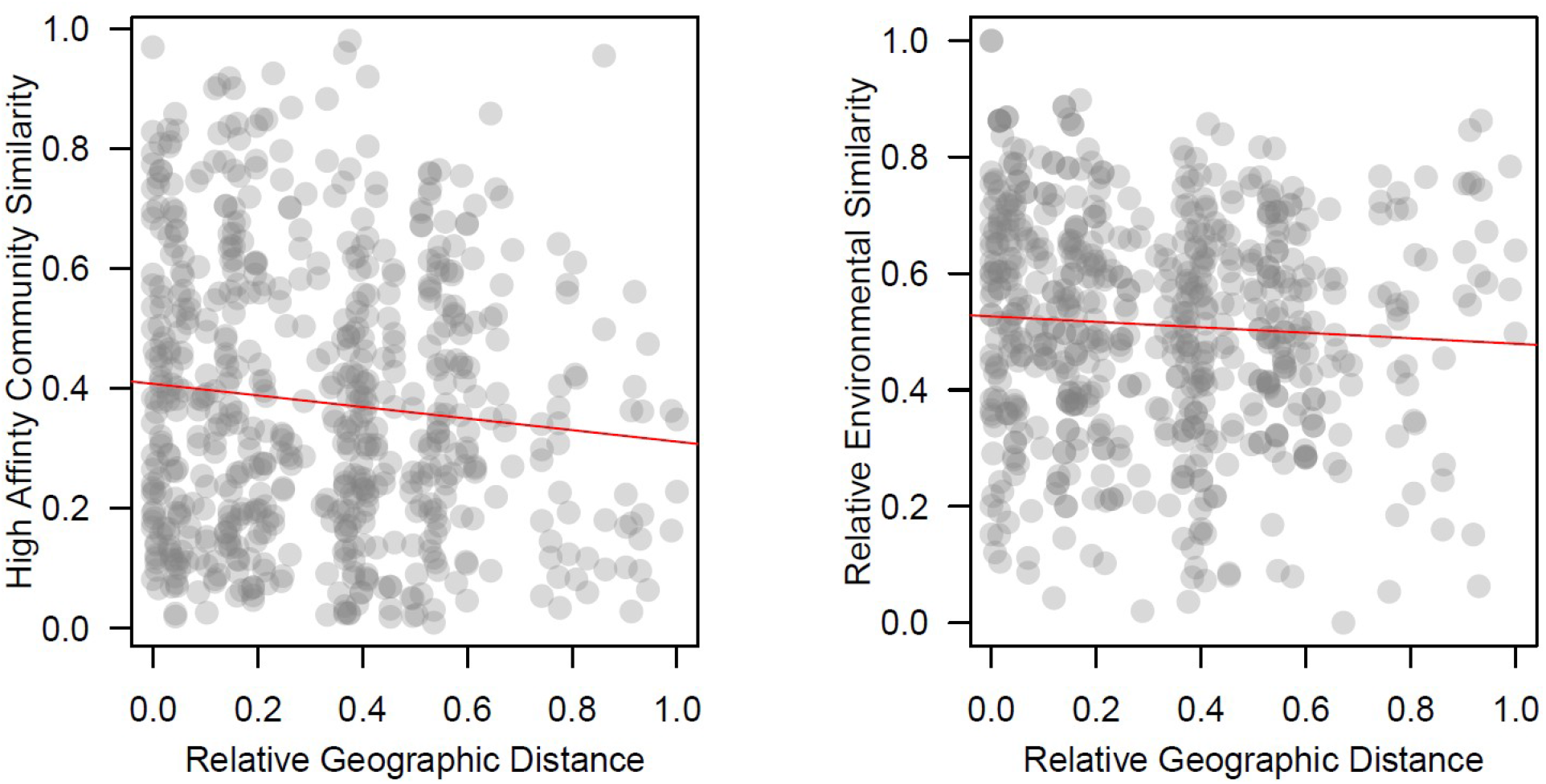
Distance decay analyses of the high affinity methanotrophs. The change in community similarity with geographic distance was statistically indistinguishable from the change in environmental similarity with distance suggesting the organisms are adept at dispersing through the environment. Note that the samples from Cave 38 have been removed from this analysis due to the large geographic separation of these samples from the rest of the samples.

### Relationships with cave air CH_**4**_ **concentrations**

The CH_4_ concentration in cave air from caves in the USA was positively correlated with the relative abundance of methanotrophs (Figure 5, Spearman’s test, *ρ* = 0.40, S = 5451, *P* < 0.01). This relationship was largely driven by high-affinity methanotrophs (Spearman’s test, *ρ* = 0.42, S = 5778, *P* < 0.01). The relative abundances of low-affinity methanotrophs were not correlated with the CH_4_ concentration (Low-affinity methanotrophs: Spearman’s test, *ρ* = 0.03, S = 9601, *P* = 0.86). The highest relative abundance of methanotrophs at any site occurs in a sample from Cave 38 which has internal sources of CH_4_ and hydrogen sulfide (H_2_S) (27), however samples from this cave also contained low relative abundances of methanotrophs.

**Figure 5:**
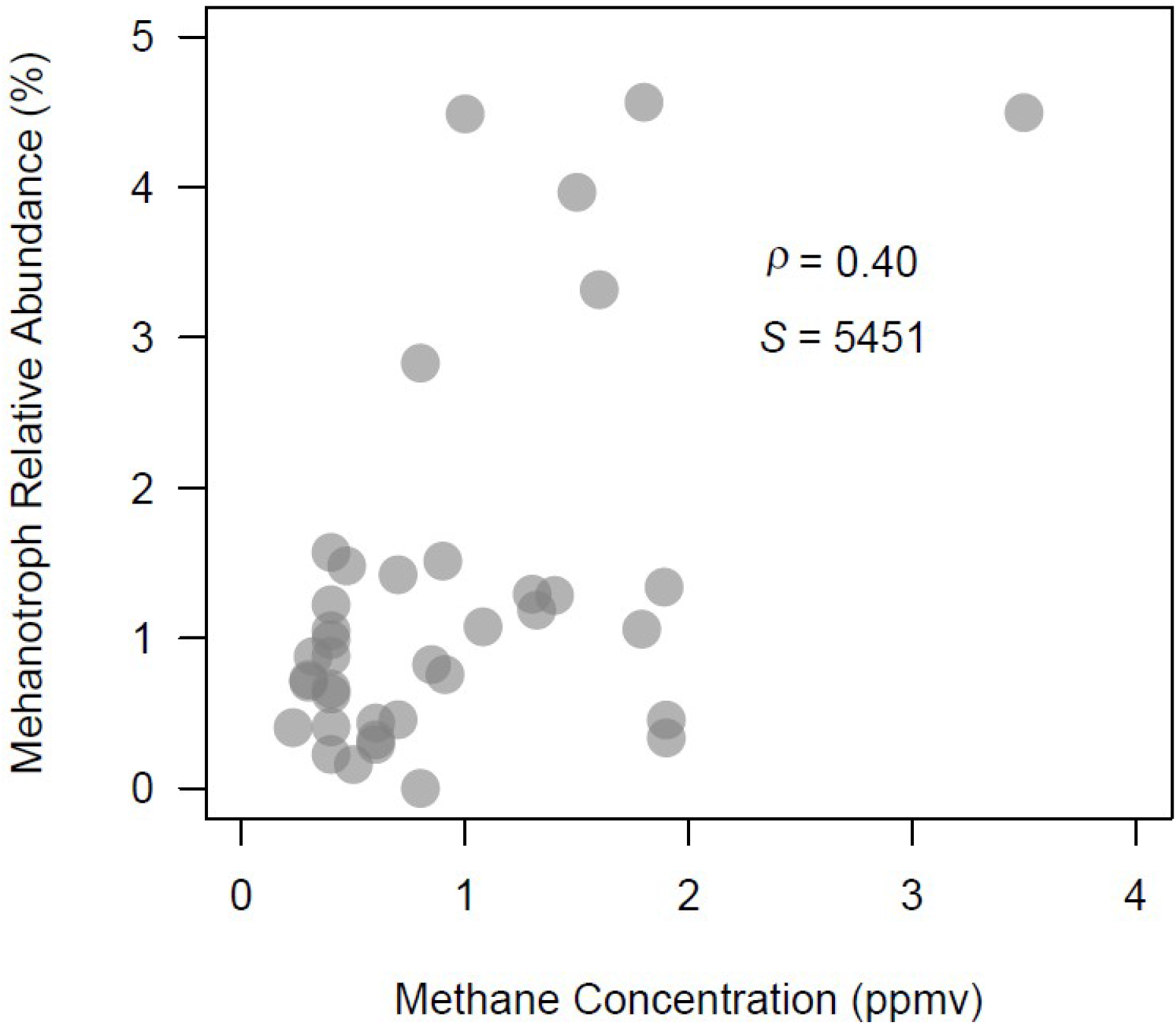
The relative abundance of members of the methanotrophic community plotted against the CH_4_ concentration at each sample location (*ρ* = 0.40, S-statistic: S = 5451, *p* < 0.01).

## DISCUSSION

### Methanotroph community composition and diversity in caves

Methanotrophs are widespread in cave environments, with high-affinity methanotrophs dominating these ecosystems. The dominance of methanotrophs from the high-affinity clades in the sampled caves is consistent with observations that have been made in other low CH_4_ environments. For example, high-affinity taxa dominate methanotroph assemblages in forest soils, Arctic soils, and soils associated with basaltic rocks (4, 27, 28). Additionally, the high relative abundance of methanotrophs in the cave soils from this study, and in particular of the USC-*γ* clade, agrees with observations from other caves where relative abundances of high-affinity methanotrophs can exceed 10 % of the entire bacterial community (8, 12, 14, 15). The relative abundances of methanotrophs observed in our study are higher than the observations of methanotrophs from caves in Vietnam (7) and contrasts with the relatively depauperate assemblage found in a Spanish cave, which reportedly consisted of only three species (29). The highest relative abundance of methanotrophs at any site occurs in a sample from Cave 38 which has an internal CH_4_ source (30).

Although less abundant, low-affinity methanotrophs from the Alphaproteobacteria and Gammproteobacteria were also commonly observed in our cave survey. Low-affinity alphaproteobacterial methanotrophs are typically more abundant in aerated soils where they have access to oxygen, whereas low-affinity gammaproteobacterial methanotrophs are more often found in environments with high CH_4_ concentrations and low oxygen availability (17, 18). Cave soils are typically nutrient poor (31, 32), implying that anoxic conditions created by the decay of organic matter are not likely to be widespread. Thus, the presence of low-affinity gammaproteobacterial methanotrophs in cave soils may be limited to local pockets of anoxia which have been suggested based on observations of limited methanogenesis in cave systems (10, 11).

We identified a core group of three OTUs that were present in the studied caves. This core community was derived entirely of high-affinity methanotrophs, suggesting that these taxa are well attuned to subterranean environments that they are able to disperse between cave ecosystems. This interpretation is consistent with previous observations suggesting that high-affinity methanotrophs have high-dispersal capabilities. For example, methanotrophs from a newly formed soil in Hawai’i showed close taxonomic affiliations with Arctic methanotrophs (4). Given that microbes are known to travel intercontinental distances on windblown dust (33), it is possible that surface dwelling high-affinity methanotrophs (27, 34) may be carried into caves on windblown dust or other passive vectors.

It is important to note that our survey presents a fairly conservative estimate for diversity and abundance of methanotrophs in cave ecosystems. In the bacteria, methanotrophy is carried out through the presence of the enzyme methane monooxygenase. Methanotrophs are viewed as specialists with regard to their metabolism (35), and, as a result, once the enzymatic machinery has been established in a lineage, it is likely to be conserved since there are barriers to evolving back into generalist species (36). Recent findings suggest that some high-affinity methanotrophs may be able to use other trace gases for carbon and energy, however (37). Often, diversity inventories of methanotrophs focus on sequences of the alpha subunit of the particulate methane monooxygenase enzyme (34). 16S rRNA analysis of methanotrophs may therefore miss some individuals in a community. However, in some studies, it has been shown that the two approaches yield similar estimates of diversity (38).

### Microenvironmental influences on methanotroph communities

Our results indicate that soil properties can play an important role in structuring methanotroph communities. The relationship between soil mineralogy and methanotrophs is most likely related to the abundance of trace metals. For example, copper is the trace metal responsible for the oxidation of CH_4_ in particulate methane monooxygenase enzyme. The methanotroph community also showed relationships with the abundance of clinochlorite, muscovite, and microcline. These minerals are associated with felsic metamorphic and igneous rocks, and metals like copper are associated with these kinds of rocks as well (39).

In contrast, the grain size of the cave soils did not influence the abundance or composition of methanotrophs, even though these edaphic characteristics strongly affect the physical and chemical properties that influence metabolism. Previous research observed that methanotrophs tend to be more abundant in silt and clay fractions of soils because these fractions made up the bulk of the soils themselves (40, 41). Despite some previous observations showing relationships between methanotrophs and soil characteristics, other studies have also shown absences of relationships between methanotroph community composition and soil characteristics (42). The importance of soil grain size in structuring methanotroph communities remains an open question.

### Biogeography of methanotroph communities

We observed regional changes in methanotrophic communities in caves. The community composition of high-affinity methanotrophs changed at the same rate as measures of environmental similarity compared to changes in the distance between the samples. This suggests that process like mass effects— which are dominant when organisms are able to actively disperse through their environment— are operating on this community (43). This observation aligns with past work indicating that high affinity methanotrophs are able to disperse through the environment (4), and the observation that three OTU accounted for 97 % of the total methanotrophic community. Dispersal through surface soils (27, 34) to caves in drip waters, and transport into caves on windblown dust (30) represent two possible ways for methanotrophs to disperse into cave systems.

The community composition of low-affinity methanotrophs did not change appreciably with geographic distance nor did measures of environmental similarity for these samples. This suggests that neutral processes are operating on these communities (43) and that random birth-death processes, or stochastic dispersal event are important in determining the distribution and abundance of methanotroph popoulations. These processes are thought to operate more strongly on communities that are not strongly influenced by the environment and are the strongest in small populations (44). This suggests that these communities may be being passively dispersed into the caves.

Biogeographic patterns at the local scale were also relatively weak. The composition and relative abundance of methanotrophic communities at the entrance versus the interior of caves were statistically indistinguishable. This suggests that environmental conditions at the entrance and interior of the cave are similar in ways the select for similar methanotroph communities. Alternatively, similarity of methanotrophs at the relatively small spatial scale may reflect repeated dispersal events in and out of the cave owing to air exchange that accompanies temperature fluctuations over daily and seasonal periods. Cave size and shape may also contribute to the exchange of microbial propagules between above ground and subterranean habitats.

### Methanotroph relative abundance and cave air CH_4_ concentrations

The relative abundance of methanotrophs was positively correlated with CH_4_ concentrations in cave air. This pattern was driven by the high-affinity methanotrophs. High-affinity methanotrophs are thought to derive a significant portion of their energy from CH_4_ at atmospheric concentrations and less energy is available for these organisms at locations in caves with sub-atmospheric CH_4_ concentrations. Some of the variability observed in the relationship between high-affinity methanotrophs and cave-air CH_4_ concentrations may be explained by daily and seasonal variation in cave air flow patterns. Many caves are known to exhibit faster air flow when the external atmospheric temperature is lower than the temperature of cave air. This leads to higher CH_4_ concentrations in caves during the winter (10, 45). One result of this is that the measured CH_4_ concentration at a particular point in a cave may or may not be representative of the CH_4_ flux at that location, and the CH_4_ flux at a particular site is likely a strong driver of methanotroph relative abundance.

## CONCLUSIONS

The structure of cave methanotroph communities appears to be best understood in terms of how cave methanotrophs access CH_4_. Cave methanotrophic communities showed relationships with cave-air CH_4_ concentrations. Additionally, high-affinity methanotrophs were numerically dominant. This community structure mirrors that of other cave communities, and other methanotroph communities that appear to consume atmospheric CH_4_ (8, 12, 14, 15, 38). High-affinity methanotrophs also comprised a core methanotrophic community with three members present together in 79 % of samples. Cave methanotroph communities showed relationships to minerals in the environment that are known to be related to the abundance of Cu, an essential trace metal for methane oxidation. These lines of evidence suggest these communities are actively consuming CH_4_ within the cave environment. The cave methanotrophic community assemblage appears to be actively dispersing across cave environments, suggesting that the dispersal barrier for high-affinity methanotrophs from cave to cave may be low. Questions remain about the relative abundance of methanotrophs in cave communities and the CH_4_ flux at a particular site. Further research may be inclined to investigate how USC-*γ* and USC-α methanotrophs partition themselves across low CH_4_ availability niches.

## MATERIALS AND METHODS

### Data and software availability

This Targeted Locus Study project has been deposited at DDBJ/EMBL/GenBank under the accession KCRG00000000. The version described in this paper is the first version, KCRG00000000.1. All code and data used in this study can be found in a public GitHub repository (https://github.com/websterkgd/CaveMethanotrophs).

### Microbial sampling

We sampled microbial communities from limestone caves along the western front of the Appalachian fold and thrust belt, in intracratonic settings of the USA, and in the Sierra Madre of Mexico. We obtained 42 cave soil samples along transects from their entrances to interiors (Data available at: https://github.com/websterkgd/CaveMethanotrophs). Cave soil samples were only collected from locations that had accumulations of sediments and included walls and floors. Samples were collected using a spatula that had been sterilized with 70 volume % ethanol in water from locations that were 0.1 m^2^ in area (46). Samples were stored on ice until they could be transferred to a −80 °C freezer.

### Environmental variables

We measured multiple environmental variables to assess factors that potentially influence the composition of cave methanotrophic communities. We measured CH_4_ concentrations *in-situ* with Fourier Transform Infrared spectroscopy (FTIR) (Gasmet DX4030 – Milton Keynes, United Kingdom) or we collected discrete air samples collected in the field for measurement the laboratory with FTIR spectroscopy or gas-chromatography (Varian – Agilent Technologies, Palo Alto, California). Some of the CH_4_ concentrations listed in this study have been previously published (11, 30), and new data were collected according to methods reported in the same publications. In cases where cave maps were available, the distance from the nearest entrance to the sampling location was calculated along the length of the cave passages. We measured cave soil grain-size distributions with a Malvern Mastersizer 3000 (Malvern Instruments Inc., Westborough Massachusetts). Raw data from the Mastersizer were converted to % gravel, % sand, % silt, and % clay sized particles by volume using the GRADISTAT software package (47). Data used for statistical analyses are available in a public GitHub repository (https://github.com/websterkgd/CaveMethanotrophs). Latitude and longitude information, however, have been removed from the data to protect the individual caves. This information can be obtained by contacting the authors directly.

### Cave soil mineralogy

We randomly selected 14 samples for mineralogical analysis using X-ray powder diffraction. Soil sediments were powdered to < 5 μm using a mortar, pestle, and acetone. Following the methods of Furmann et al. (48), we used Bruker D8 Advance X-ray diffractometer with a Sol-X solid-state detector and a W X-ray tube operated at 30 mA and 40 kV to identify mineral phases. We placed cave sediment powders in an aluminum large frontpacked mount. Packed mounts were scanned from 2° to 70° using a count time of 2 s per 0.02° step. Rietveld refinements were used to determine the abundances of the minerals present in the sample and were quantified with TOPAS software. Proportional abundances of the minerals from the cave soils are reported in Table 1.

**Table 1:** The proportional abundance of minerials from the sampled locations

| Sample | Cave | Quartz | Muscovite | Clinochlorite | Albite | Orthoclase | Calcite | Microcline | Dolomite | Anorthite | Gypsum |
| --- | --- | --- | --- | --- | --- | --- | --- | --- | --- | --- | --- |
| 21-10d | Cave 21 | 0.72 | 0.14 | 0.01 | 0.06 | 0.04 | 0.00 | 0.00 | 0.00 | 0.03 | 0.00 |
| 22-11d | Cave 22 | 0.65 | 0.16 | 0.01 | 0.07 | 0.00 | 0.01 | 0.11 | 0.00 | 0.00 | 0.00 |
| 17-1b | Cave 17 | 0.48 | 0.28 | 0.01 | 0.00 | 0.00 | 0.23 | 0.00 | 0.00 | 0.00 | 0.00 |
| 17-1c | Cave 17 | 0.89 | 0.09 | 0.00 | 0.00 | 0.00 | 0.03 | 0.00 | 0.00 | 0.00 | 0.00 |
| 27-5b | Cave 27 | 0.33 | 0.19 | 0.01 | 0.06 | 0.00 | 0.07 | 0.28 | 0.05 | 0.00 | 0.00 |
| 27-5d | Cave 27 | 0.50 | 0.15 | 0.01 | 0.00 | 0.00 | 0.16 | 0.14 | 0.04 | 0.00 | 0.00 |
| 30-8d | Cave 30 | 0.40 | 0.13 | 0.00 | 0.02 | 0.00 | 0.19 | 0.09 | 0.18 | 0.00 | 0.00 |
| 31-9b | Cave 31 | 0.40 | 0.20 | 0.01 | 0.00 | 0.00 | 0.05 | 0.16 | 0.19 | 0.00 | 0.00 |
| 31-9c | Cave 31 | 0.14 | 0.20 | 0.00 | 0.00 | 0.00 | 0.06 | 0.08 | 0.52 | 0.00 | 0.00 |
| 35-1 | Cave 35 | 0.70 | 0.09 | 0.00 | 0.08 | 0.04 | 0.00 | 0.08 | 0.00 | 0.00 | 0.00 |
| 38-9 | Cave 38 | 0.38 | 0.29 | 0.00 | 0.00 | 0.00 | 0.00 | 0.00 | 0.00 | 0.00 | 0.33 |
| 10-1 | Cave 10 | 0.76 | 0.11 | 0.00 | 0.06 | 0.00 | 0.00 | 0.06 | 0.01 | 0.00 | 0.00 |
| 4-1 | Cave 4 | 0.21 | 0.24 | 0.00 | 0.00 | 0.00 | 0.55 | 0.00 | 0.00 | 0.00 | 0.00 |
| 2-3 | Cave 2 | 0.82 | 0.11 | 0.00 | 0.00 | 0.00 | 0.00 | 0.07 | 0.00 | 0.00 | 0.00 |
| 2-2 | Cave 2 | 0.75 | 0.12 | 0.00 | 0.04 | 0.00 | 0.02 | 0.07 | 0.00 | 0.00 | 0.00 |
| 2-1 | Cave 2 | 0.74 | 0.10 | 0.00 | 0.06 | 0.00 | 0.00 | 0.10 | 0.00 | 0.00 | 0.00 |

### Molecular techniques

We extracted genomic DNA from cave soil samples with a MoBio PowerSoilTM extraction kit (MoBio, Carlsbad, California USA). About 10 ng of extracted DNA was used as a template for amplification by polymerase chain reaction (PCR). The V4 region of the 16S rRNA gene was amplified using 515F and 806R with barcoded primers designed to work with the Illumina MiSeq platform (49). PCR reactions were as follows: 3 min denaturing step at 94 °C, followed by 30 cycles of 94 °C for 45 s, 50 °C for 60 s, and 72 °C for 90 s. A final 10-minute extension was carried out at 72 °C. Quality of the PCR products was checked by gel electrophoresis. Amplified DNA was cleaned using a commercial kit (Beckman Coulter Agencourt AMPure XP, Indianapolis, Indiana, USA) before quantification using the QuanIt PicoGreen kit (Invitrogen) and after which aliquots were pooled at a final concentration of 20 ng per sample.

We sequenced the pooled sample containing PCR products using Illumina MiSeq technology (Illumina Reagent Kit v2, 500 reaction kit) at the Center for Bioinformatics and Genomics at Indiana University. Data quality and unique sequences obtained from the PCR amplifications were analyzed using mothur (50). DNA sequence data were aligned using the Needleman algorithm and read lengths were limited to 197 base pairs. Sequences matching chimeras were removed using VSEARCH (51). Sequences that matched chloroplasts, Archaea, and other non-bacterial sequences were also removed. OTUs were created by binning the data at 97 % sequence similarity using the opticlust algorithm (52). OTUs were identified using the SILVA reference database (version 132).

### Classification of methanotrophs

We identified methanotrophs from the recovered 16S rRNA sequences using a three step process. First, we compiled a database of organisms known methanotrophs through literature surveys. This included assembling lists of low-affinity methanotrophs from the Methylococcaceae, Methylocystaceae, and Beijernickiaceae (34, 53). We used the genera *Methylacidiphilum, Methylacidimicrobium* to identify methanotrophs belonging to the Verrucomicrobia (34, 54). High-affinity methanotrophs were defined as organisms belonging to the USC-*α* and -*γ* clades, and *Methylocapsa gorgona* MG08 (13, 34, 53). Second, organisms from this database were checked against the identifications assigned to the OTUs during the initial analysis from the SILVA database (version 132). Third, we compiled a FASTA file of the 16S rRNA gene of 71 known methanotrophs. 16S rRNA sequences for USC-*α* and USC-*γ* methanotrophs were obtained using published 16S rRNA sequences for USC-*α* from (12), for USC-*γ* from (13), and for *Methylocapsa gorgona* MG08 from (37).. This file was compared against the FASTA file of all organisms in the dataset using the *usearch_global* command to identify putative methanotrophs that may have been missed through the SILVA analysis (usearch v.11.0.667) (55). This list of putative methanotophs was then compared against the NCBI database. If the sequences did not match an organism capable of the function of interest at 98 % similarity they were excluded from downstream analysis. The process just described was intended to prevent the false classification of an organism as a methanotroph. The compiled methanotroph FASTA file used in our analysis is available in a public GitHub repository (https://github.com/websterkgd/CaveMethanotrophs).

### Statistical analyses

We assessed the relationships between methanotrophs and environmental conditions through a series of statistical analyses. First, to avoid complication arising from unavoidable variation in the total number of reads between samples, we calculated the relative abundance for each taxon within a sample. Next, to quantify among cave variation, we calculated pairwise dissimilarities using the Bray-Curtis metric. The core methanotroph community was characterized by including taxa that were found in 60 % or more of our samples. We used this cutoff because it is in the middle of the range of what has been used in other studies (23, 24). We evaluated the effect of sediment mineralogy and grain size on the multivariate composition of methanotroph communities using canonical correspondence analysis (CCA) with the *vegan* package in R (56). If a sample did not have associated mineralogical data, it was not included in the CCA analysis.

To gain insight into the biogeography of cave methanotrophs, we examined distance decay relationships (DDR) while also comparing community composition along gradients from cave exteriors to cave interiors. We analyzed DDR by plotting Bray-Curtis similarity of methanotroph composition against environmental and geographic distance calculated as standardized Euclidean distances. Samples from Cave 38 (Cueva de Villa Luz) were excluded in DDR analyses since they were roughly 2000 km away from the nearest sample. Slopes for geographic and environmental DDRs were calculated using ordinary least squares (23). We then tested for differences in the slopes using the statistical test for a difference of slope between regression lines (57). We used Spearman’s test to examine how methanotrophic communities changed along gradients from cave exteriors to interiors. Finally, we tested for a relationship between methanotroph relative abundance and CH_4_ concentrations using Spearman’s test.

## ACKNOWLEDGMENTS

This material is based upon work supported by the U.S. Department of Energy, Office of Science, Office of Basic Energy Sciences under Award Number DE-SC0006978 to AS. Additional support was provided by a National Speleological Society research grant awarded to KDW, the National Geographic Society Expeditions Council (Grant #EC0644-13 to PJB), the National Science Foundation (1442246 to JTL), the US Army Research Office (W911NF-14-1-0411 to JTL) and the National Aeronautics and Space Administration (80NSSC20K0618 to JTL). We thank Brent Lehmkuhl for assistance in the laboratory. Members of the Bloomington Indiana Grotto facilitated access to caves in Indiana and Kentucky. We thank Rodolfo Gómez Cruz for his support at Cueva de Villa Luz (Cave 38). ML Larsen, MM Muscarella, and KJ Locey aided in data analysis and interpretation. KDW thanks A Barberán for discussions on genetic analysis that resulted in development of the methods used in this paper. KDW thanks the Planetary Science Institute Mars Laboratory for providing computing resources. The authors declare no conflicts of interest.

## Notes

https://github.com/websterkgd/CaveMethanotrophs

### Competing Interest Statement

The authors have declared no competing interest.

### Summary of Updates

This revision reflects a redone analysis.

